# Miniscule differences between the sex chromosomes in the giant genome of a salamander, *Ambystoma mexicanum*

**DOI:** 10.1101/354092

**Authors:** Melissa C. Keinath, Nataliya Timoshevskaya, Vladimir A. Timoshevskiy, S. Randal Voss, Jeramiah J. Smith

## Abstract

In the Mexican axolotl (*Ambystoma mexicanum*) sex is known to be determined by a single Mendelian factor, yet the sex chromosomes of this model salamander do not exhibit morphological differentiation that is typical of many vertebrate taxa that possess a single sex-determining locus. Differentiated sex chromosomes are thought to evolve rapidly in the context of a Mendelian sex-determining gene and, therefore, undifferentiated chromosomes provide an exceptional opportunity to reconstruct early events in sex chromosome evolution. Whole chromosome sequencing, whole genome resequencing (48 individuals from a backcross of axolotl and tiger salamander) and *in situ* hybridization were used to identify a homomorphic chromosome that carries an *A. mexicanum* sex determining factor and identify sequences that are present only on the W chromosome. Altogether, these sequences cover ~300 kb, or roughly 1/100,000^th^ of the ~32 Gb genome. Notably, these W-specific sequences also contain a recently duplicated copy of the ATRX gene: a known component of mammalian sex-determining pathways. This gene (designated *ATRW*) is one of the few functional (non-repetitive) genes in the chromosomal segment and maps to the tip of chromosome 9 near the marker *E24C3*, which was previously found to be linked to the sex-determining locus. These analyses provide highly predictive markers for diagnosing sex in *A. mexicanum* and identify *ATRW* as a strong candidate for the primary sex determining locus or alternately a strong candidate for a recently acquired, sexually antagonistic gene.

**AUTHOR SUMMARY:** Sex chromosomes are thought to follow fairly stereotypical evolutionary trajectories that result in differentiation of sex-specific chromosomes. In the salamander A. mexicanum (the axolotl), sex is determined by a single Mendelian locus, yet the sex chromosomes are essentially undifferentiated, suggesting that these sex chromosomes have recently acquired a sex locus and are in the early stages of differentiating. Although Mendelian sex determination was first reported for the axolotl more than 70 years ago, no sex-specific sequences have been identified for this important model species. Here, we apply new technologies and approaches to identify and validate a tiny region of female-specific DNA within the gigantic genome of the axolotl (1/100,000th of the genome). This region contains a limited number of genes, including a duplicate copy of the ATRX gene which, has been previously shown to contribute to mammalian sex determination. Our analyses suggest that this gene, which we refer to as ATRW, evolved from a recent duplication and presents a strong candidate for the primary sex determining factor of the axolotl, or alternately a recently evolved sexually antagonistic gene.

## INTRODUCTION

In many species, sex is determined by the inheritance of highly differentiated (heteromorphic) sex chromosomes, which have evolved independently many times throughout the tree of life (1–3). Often these chromosomes differ dramatically in morphology and gene content (4–6). In mammals, males have a large, gene rich X-chromosome and a degraded, gene poor Y-chromosome, while females have two X chromosomes. In birds and many other eukaryotes, females are the heterogametic sex with a large Z and smaller W chromosome, while males are homozygous, carrying two Z chromosomes. Differentiated sex chromosomes are thought to arise through a relatively stereotypical process that begins when a sex-determining gene arises on a pair of homologous autosomes (5, 6). The acquisition of sexually antagonistic alleles, alleles that benefit one sex and are detrimental to the other, favors the fixation of mutational events that suppress recombination in the vicinity of the sex-determining locus (7, 8). Recombination suppression can lead to the accumulation of additional sexually antagonistic mutations and repetitive elements, and over time this results in the loss of nonessential parts of the Y or W chromosome, resulting in the formation of heteromorphic sex chromosomes (9).

Unlike the majority of mammals and birds with stable sex-determining systems and heteromorphic sex chromosomes, amphibians have undergone numerous evolutionary transitions between XY and ZW-type mechanisms and may possess morphologically indistinguishable (homomorphic) sex chromosomes, like those of the axolotl (10–13). Homomorphic sex chromosomes are not altogether rare among animals, with examples in fish (14), birds (15), reptiles (16) and amphibians (17). Among most amphibians that have been investigated, homomorphy is prevalent (17–19). It has been suggested that a majority of salamanders have homomorphic sex chromosomes (18, 20), however, evidence for genetic sex determination in most species is largely based on the observation of 1:1 sex ratios from clutches without thorough demonstration of Mendelian inheritance.

Early developmental/genetic experiments revealed a ZW type sex-determining mechanism for *A. mexicanum* (21–23). The first experiment to test for female heterogamety involved sex reversal through implantation of a testis preprimordium from a donor embryo to a host female embryo. The prospective ovary developed instead into a functional testis. This sex-reversed male was then crossed with a normal female (24). It was expected that if the female were homozygous for sex (XX), the offspring would all be female. If the female were heterozygous for sex (ZW), however, the offspring would have an approximate female to male ratio of 3:1. Two matings with the sex-reversed animals produced a combined 26.1% males, consistent with the hypothesis that the male was indeed a sex-reversed female with ZW chromosomes (21, 24). Subsequent studies showed normal sex ratios from matings with the F1 males and most of the F1 females, but several of the F1 females produced spawns of all females, suggesting they carried the unusual WW genotype (24).

Following these foundational studies, early genetic mapping studies used cold shock to inhibit meiosis II and assessed triploid phenotypes to estimate the frequencies of equatorial separation and map distances between recessive mutations and their linked centromeres (25). Based on these analyses, the sex determining locus was predicted to occur near the end of an undefined chromosome (25) and later estimated to be 59.1 cM distal to the centromere (essentially, freely recombining) (23).

Karyotypic analyses later indicated that the smallest chromosomes were heteromorphic in *Ambystoma* species, suggesting that the smallest pair of chromosomes carried the Mendelian sex determining factor in *A. mexicanum* (26) and in the *A. jeffersonianum* species complex (27). However, more recent linkage mapping studies indicated that sex was determined by a locus on one of the larger linkage groups (26, 28), and chromosome sequencing studies have demonstrated that the smallest chromosomes do not carry the sex determining region (29, 30). Notably, extensive cytogenetic studies performed by Callan (31), including the use of cold treatments to add constrictions to chromosomes and examination of lampbrush chromosomes from developing oocytes, revealed no features that could be associated with differentiated sex chromosomes. These analyses not only indicated that the sex chromosomes were apparently identical to one another, but also revealed that mitotic chromosomes 9, 10 and 11 were essentially indistinguishable from one another (31).

More recently, meiotic mapping of polymorphisms within controlled crosses localized the sex-determining region to the tip of *Ambystoma* LG9 (previously designated LG5) distal to the marker *E24C3* (29). These crosses included a mapping panel that was generated by backcrossing female *A. mexicanum/A. tigrinum* hybrids with male *A. mexicanum*. These crosses also revealed no difference in recombination frequencies between the sexes. However, these studies were somewhat limited by the fact that they did not sample large numbers of markers in close proximity to the sex locus or W-specific sequences (29). Taken together, analyses of the *Ambystoma* sex determination suggest that the sex chromosomes are largely undifferentiated and that, presumably, the sex chromosomes arose recently within the tiger salamander species complex.

To identify sex-linked (W-specific) regions in the undifferentiated sex chromosomes of axolotl, we generated sequence reads for 48 individuals of known sex that were derived from a backcross (*A. mexicanum/A. tigrinum* X *A. mexicanum*). These reads were then aligned to an existing reference genome from a female axolotl (30, 32) (www.ambystoma.org). Analyses of read coverage identified 152 putative W-linked sequences, including two genes, an *ATRX* paralog and an ortholog of *MAP2K3*. The W-linked ATRX paralog, *ATRW*, is estimated to have duplicated within the last 20 million years, providing an estimate of the possible origin of the sex-determining locus in the tiger salamander species complex. In addition, we anticipate that these sex-linked markers will be useful for identifying sex in juvenile axolotls within lab-reared populations, where sex is an important covariate for experimental studies, including studies of metamorphosis and regeneration (28, 33).

## RESULTS

### Identification of the sex-bearing chromosomes by FISH

Previous studies have demonstrated that sex is linked to the marker *E24C3*, at a distance of ~5.9 cM distal to the terminal marker on LG9 (29). Consistent with linkage analyses, *E24C3* was detected near the tip of an average-sized chromosome (Figure 1). A second BAC corresponding to a marker from the opposite end of LG9 (*E12A6*) localized to the opposite tip of the same chromosome, indicating that this chromosome corresponds precisely to LG9 (Figure 1). Notably, the BAC carrying *E12A6* also cross-hybridized with the centromere of all chromosomes, a feature that could potentially be useful in estimating distances of genes to their respective centromeres.

**Figure 1.**
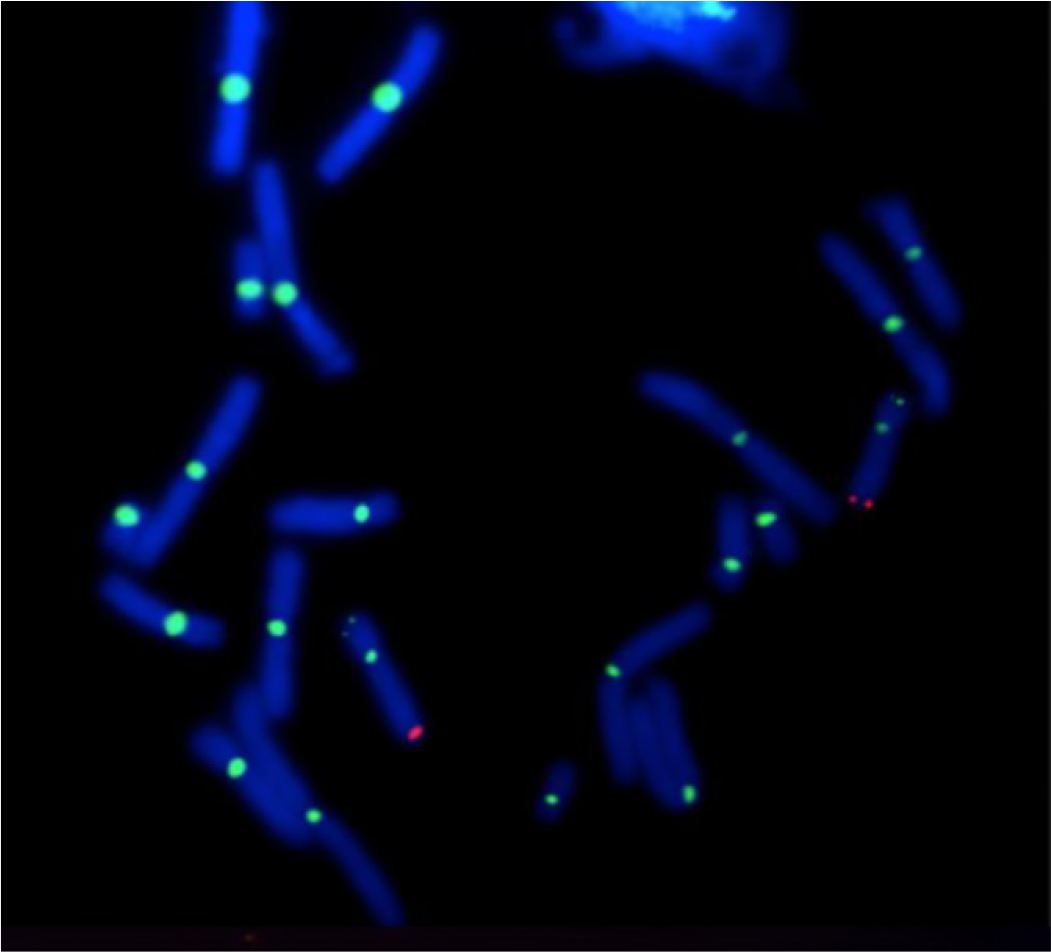
FISH of sex-linked BACs. FISH localizes two markers (*E24C3* and *E12A6*) associated with the sex locus, *ambysex*, on a DAPI stained metaphase spread of chromosomes from an axolotl embryo of unknown sex. *E24C3* is labeled with cy3 (red) and *E12A6* is labeled with fluorescein (green).

### Laser capture, sequencing and assembly of the Z chromosome

In an attempt to increase the number of markers that could be associated with the sex chromosome, we performed laser-capture sequencing on a chromosome corresponding to LG9. This library was generated from a single dyad that was collected in a larger series of studies on laser capture microscopy of axolotl chromosomes (34). The sex chromosome library contained a total of ~143 M reads between 40 and 100 bp after trimming and contained 995 reads that mapped to 23 distinct markers (transcripts) that had been previously placed on LG9 (Figure 2). In total, this initial sequencing run accounted for 40% of the markers that are known to exist on the linkage group, which was considered strong evidence that this library sampled the sex chromosome. Given this support, an additional lane of sequencing was performed, yielding ~936 M additional reads (for a total of 1,078,893,614 reads). After trimming, ~542 M reads remained. Alignment to human and bacterial genomes revealed that 1.7% and 0.1% of trimmed reads aligned concordantly to the human genome and bacterial genomes, respectively. These were considered contaminants and were removed from subsequent analyses. Of the remaining reads, 68,844 aligned to 40 LG9 contigs representing 70% of the known markers on LG9 (Figure 2). An error-corrected assembly of these data yielded a total of 1,232,131 scaffolds totaling 242.4 Mb with a scaffold N50 length of 295 bp, and contig N50 length of 126 bp. (Table 1: results from other chromosomes are shown for comparison purposes). We also used this library to identify a set of scaffolds from a recently published assembly of a male axolotl genome that could be assigned to the Z chromosome on the basis of sequence coverage. This analysis yielded 2531 scaffolds spanning a total of 1.02 Gb (Supplementary Table 1).

**Table 1.** Summary statistics for LG9, AM13 and AM14 chromosome assemblies. Summary statistics for de novo assembly of sequence data from the sex chromosome, which corresponds to linkage group 9 (LG9) as well as AM13 and AM14 for comparison as previously published (30). Chromosomes 13 and 14 correspond to *A. mexicanum* linkage groups 15/17 (LG15/17) and linkage group 14 (LG14), respectively. Statistics are presented for assemblies of raw sequence data (R) and assemblies post error correction (EC).

|  | Assembly |  |  | Contig |  | Scaffold |  |
| --- | --- | --- | --- | --- | --- | --- | --- |
|  | Length<br>(Mb) | Number<br>of<br>Scaffolds | Number<br>of<br>Singletons | N50 Length<br>Improvement | Proportion<br>Scaffolded | N50 Length<br>Improvement | Number<br>>N50 |
| LG9 (R) | 189.7 | 1,054,224 | 760,174 | 118 | 0.352 | 256 | 285,628 |
| LG9<br>(EC) | 242.4 | 1,232,131 | 866,817 | 126 (6.8%) | 0.429 | 295(15.2%) | 335,062 |
| LG15/17<br>(R) | 302.5 | 604,617 | 243,354 | 231 | 0.598 | 705 | 136,682 |
| LG15/17<br>(EC) | 210.9 | 353,381 | 126,169 | 295 (28%) | 0.643 | 830 (18%) | 82,835 |
| LG14<br>(R) | 180.4 | 367,575 | 145,951 | 232 | 0.603 | 686 | 83,979 |
| LG14<br>(EC) | 143.0 | 258,214 | 93,931 | 290 (25%) | 0.636 | 765 (12%) | 62,022 |

**Figure 2.**
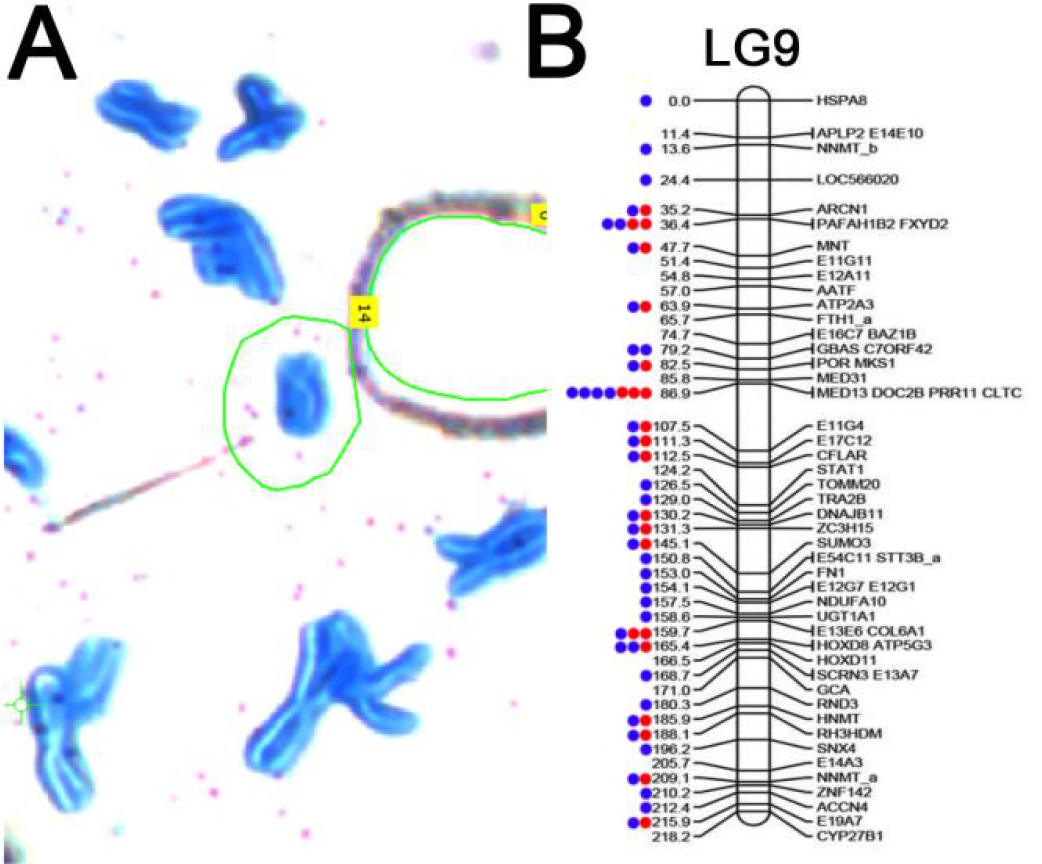
Individual sex chromosome dyad alignment results on LG9. Read mapping was used to assess the specificity of the laser capture, amplified library of the sex chromosome dyad. **A)** A partial metaphase spread of axolotl chromosomes stained with Giemsa on a membrane slide. The sex chromosome is circled in green. **B)** The distribution of markers sampled from the sex chromosome (LG9) via targeted sequencing of individual chromosomes. Dots represent markers with mapped reads from a single library. Red denotes the first sequencing attempt using the DNA-seq kit with 48 total barcoded samples on a single lane of an Illumina HiSeq flowcell. Blue denotes re-sequencing of the same chromosome library on a single lane.

Alignments between the sex chromosome assembly and *Ambystoma* reference transcripts (www.ambystoma.org) were used to identify genes that are encoded on the sex chromosome. These genes were aligned to the chicken genome assembly to confirm that homologs from the axolotl sex chromosome were heavily enriched on chicken chromosomes 7, 19 and 24, and similar enrichment was observed among scaffolds assigned to the Z from the male assembly, consistent with previous findings (Figure 3A, Supplementary Table 1) (35). Alignments to the newt (*Notophthalmus viridescens*) linkage map support previous analyses demonstrating that axolotl LG9 is homologous to newt LG7 (36), revealing strong conservation for the sex chromosome over the last 150 million years (Figure 3B). While a ZW-type mechanism for sex determination has been inferred for the newt (37), the exact chromosome that determines sex is unknown and no candidate genes currently exist.

**Figure 3.**
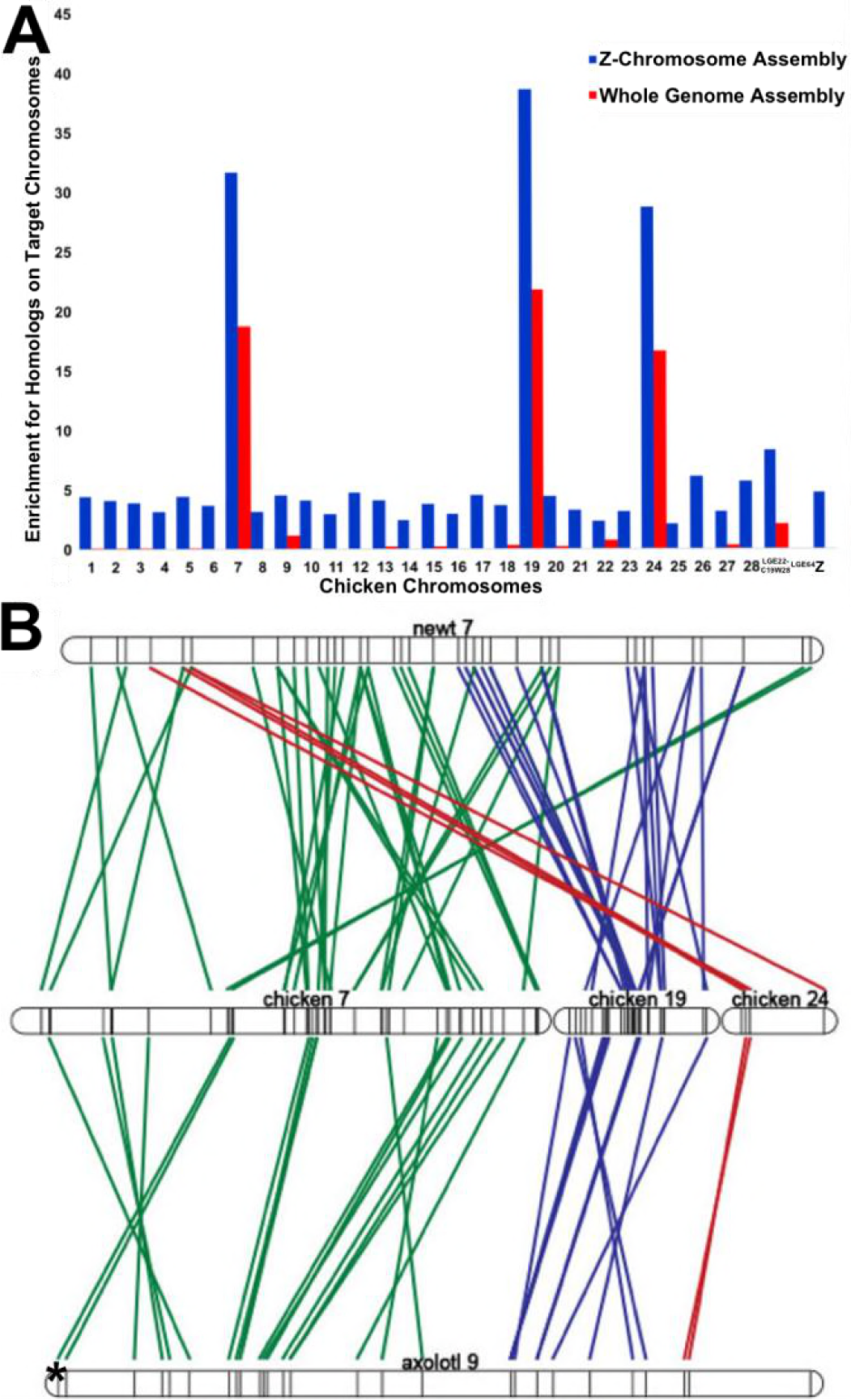
Conserved synteny for *A. mexicanum* sex chromosome. **A)** Conserved synteny between assembled A. *mexicanum* Z chromosome and the chicken genome. Tests for enrichment of Z chromosome homologs with 99% identity from read mapping-based (blue) and assembly-based (red) methods across all assembled chicken chromosomes. Enrichment scores are calculated by dividing the observed number of homologs by the total number of genes annotated to the individual chicken chromosomes (86). **B)** Conserved synteny studies show syntenic regions shared between newt linkage group 7 (top), chicken chromosomes 7,19, and 24 (middle), and axolotl LG9 (bottom). Each line corresponds to an alignment between a pair of presumptive chicken and salamander (newt or axolotl) orthologs, and the asterisk denotes the sex-specific region. Alignments involving orthologs on chicken chromosome 7 are colored green, chromosome 19 are colored blue, and chromosome 24 are red. More alignments were found between newt and chicken, as the linkage map of the newt is more dense than that of the axolotl (36).

### *In silico* identification of female-specific regions

To identify sex-specific regions of the genome, we aligned low coverage sequence data from 26 males and 22 females to both the LG9 assembly and the first public draft assembly of the axolotl genome (30, 32) (www.ambystoma.org). The draft assembly was generated using a modified version of SparseAssembler (38) from 600 Gb of HiSeq paired end reads and 640Gb of HiSeq mate pair reads. Sequencing data were produced using DNA from a female axolotl, which should contain genomic regions from both Z and W chromosomes. Notably, a recently published draft genome was generated from a male and is not expected to represent W-specific regions (39). Males and females used for re-sequencing efforts were drawn from a previously published meiotic mapping panel, which was used in the initial mapping of the sex locus (29). Each individual was sequenced to ~1X coverage with Illumina HiSeq short paired-end reads (125bp) resulting in ~7.4 billion total male reads and 6.4 billion total female reads. The ratio of female to male coverage was calculated across ~10.5M intervals covering ~19 Gb of the draft assembly. Genome-wide coverage ratios generally fell within a tight distribution centered on equal coverage, after accounting for initial differences in average depth of coverage (Figure 4). Intervals were considered to be candidate sex-specific regions if enrichment scores [log_2_ (female coverage/adjusted male coverage)] exceeded two. In total, these analyses identified only 201 candidate female-specific intervals that were contained within 109 genomic scaffolds, with 20 genomic scaffolds having 2 or more intervals (Supplementary Table 2). The combined size of these intervals is approximately 300Kb or ~0.0094% of the genome. 47 intervals were represented by zero male reads, and the average male coverage of male reads for other intervals ranged from 0.002 to 8.63.

**Figure 4.**
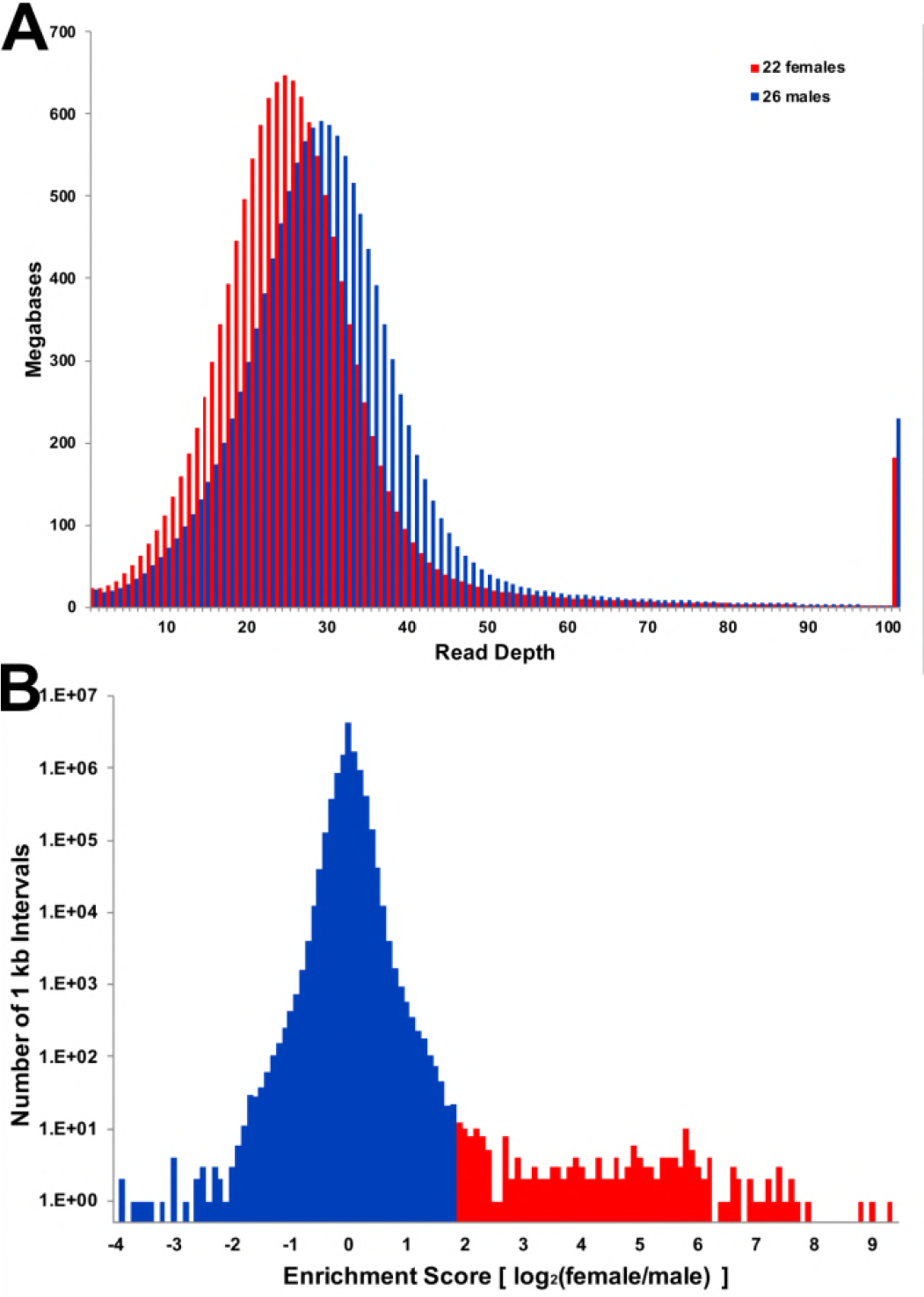
Distribution of read depth from combined female and males sequencing data. **A)** Sequence reads from 48 individuals were mapped separately to the female whole genome assembly, then alignment files were merged across all individuals of a given sex (22 females and 26 males). Values represent the number of base pairs of the reference assembly that were sampled at a given depth of coverage. These distributions reveal that the modal coverage of reads from females was lower than the coverage of males, ~25X and ~29X, respectively, consistent with random sampling of sequences across individuals. There is no overtly visible evidence that female sequences map to a larger proportion of the approximate single copy sequence within the female genome. **B)** The distribution of coverage ratios is tightly centered on equal coverage and only a small tail corresponds to intervals with higher sequence coverage in female relative to male.

### PCR validation of candidate regions

PCR primers were designed for all candidate scaffolds and subjected to initial PCR validation using a panel of six females and six males (Supplementary Table 3). In total, primers from 42 of the 109 scaffolds yielded specific amplicons in all females and no amplicons from males and were considered sex-specific. The combined size of these scaffolds is approximately 174Kb or ~0.0054% of the genome. Aside from the PCR validated female-specific scaffolds, primers from one scaffold were present in all females and one male, two were present in four females and no males, and four were present in a subset of the animals with no specific trend toward one sex or the other. Presumably these represent structural (insertion/deletion) variants that are segregating within the lab population of *A. mexicanum*, perhaps representing tiger salamander (*A. tigrinum*) DNA remnants that were introgressed in 1962 (40). Primers for another 46 scaffolds yielded amplification in both sexes with 14 showing brighter bands in females and two showing varying brightness across all individuals. Primers for seven other scaffolds yielded no amplification in either sex. To further investigate our PCR validation results, we retrospectively aligned predicted W-specific regions to the recently published *A. mexicanum* (male) genome. These revealed that several predicted W-specific contigs correspond to copies of repetitive elements with highly similar sequences elsewhere in the genome, which appears to explain a majority of cases wherein primers yield amplicons in both sexes or are polymorphic among males and females.

### Identifying W-specific genes

To search for evidence of sex-specific genes, all 42 validated sex-specific scaffolds were aligned (blastx) to the NCBI nonredundant protein database (41). In total, these searches yielded alignments to 17 protein-coding genes (Table 2), several of which involved weak alignments to uncharacterized proteins (N = 4) or transposable elements (N = 5). However, two scaffolds yielded strong alignments to human protein coding genes. Specifically, Scaffold SuperContig_990642 aligned to transcriptional regulator *ATRX*(*ATRX:* 65% amino acid identity) and scaffold SuperContig_1084421 aligned to mitogen-activated protein kinase kinase kinase 2-like (MAP3K2: 97% amino acid identity). Notably, a conserved syntenic ortholog of MAP3K2 would be expected to occur on LG9 and thus it seems likely that MAP3K2 resided on the ancestral LG9 sex chromosome prior to the origin of the *A. mexicanum* sex-determining locus. However, a syntenic ortholog of *ATRX* would be expected to occur within a conserved synteny on a different chromosome (LG2, LG8/12), corresponding to a large region of conservation with mammalian X chromosomes and chicken chromosome 4 (42–44).

**Table 2.** Blast results to nonredundant protein NCBI database. The table shows best match amino acid alignments for blast (blastx) (78) hit results for all 42 sex-specific scaffolds. 17 scaffolds aligned to a protein-coding gene, and most shared <40% identity. The two highest identity hits to genes were to transcriptional regulator *ATRX* by SuperContig_990642 and mitogen-activated kinase kinase kinase 2 by SuperContig_108441.

| Sex-specific<br>Scaffold | Scaffold<br>length<br>(bp) | NCBI Best Hit | Query<br>Cover | E<br>value | %<br>identity | Accession # |
| --- | --- | --- | --- | --- | --- | --- |
| SuperContig1084421 | 991 | PREDICTED: mitogen-activated protein kinase<br>kinase kinase 2-like<br>[Phaethon lepturus] | 18% | 6.00E-33 | 98% | XP_010292439.1 |
| SuperContig_990642 | 1488 | PREDICTED: transcriptional<br>regulator ATRX isoform X2<br>[Alligator sinensis] | 17% | 6.00E-13 | 64% | XP_006032758.2 |
| SuperContig_1201750 | 725 | PREDICTED:<br>uncharacterized protein<br>LOC101734340 [Xenopus<br>tropicalis] | 14% | 0.13 | 50% | XP_017945915.1 |
| SuperContig_1270996 | 631 | hypothetical protein<br>[Rhodopirellula baltica] | 12% | 9.7 | 50% | WP_011119337.1 |
| SuperContig_1240926 | 668 | PREDICTED:<br>uncharacterized protein<br>LOC106589496 [Salmo salar] | 39% | 2.00E-16 | 47% | XP_014035031.1 |
| SuperContig_481414 | 11464 | PREDICTED: dynein heavy<br>chain 11, axonemal [Xenopus<br>tropicalis] (reverse<br>transcriptase) | 5% | 1.00E-32 | 43% | XP_017952780.1 |
| SuperContig_1139773 | 843 | aminotransferase class I and<br>II [Streptomyces sp.<br>CB00455] | 17% | 6.2 | 42% | WP_073917349.1 |
| SuperContig_1398497 | 510 | hypothetical protein [Massilia sp. BSC265] | 36% | 4.1 | 40% | WP_051933638.1 |
| SuperContig_1136461 | 850 | flagellar autotomy protein [Micromonas pusilla CCMP1545] (reverse transcriptase) | 12% | 0.55 | 39% | XP_003062983.1 |
| SuperContig_1105317 | 928 | hypothetical protein A2Z37_15870 [Chloroflexi bacterium RBG_19FT_COMBO_62_14] | 12% | 6.3 | 37% | OGO67717.1 |
| SuperContig_960617 | 1857 | PREDICTED: uncharacterized protein LOC106605384 [Salmo salar] | 20% | 1.8 | 36% | XP_014056412.1 |
| SuperContig_446459 | 12684 | ORF2 protein [Salmo salar] (reverse transcriptase) | 8% | 1.00E-36 | 35% | AKP40998.1 |
| SuperContig556195 | 9021 | PREDICTED: uncharacterized protein LOC108708171 [Xenopus laevis] | 19% | 2.00E-70 | 34% | XP_018102087.1 |
| SuperContig_981147 | 1581 | PREDICTED: LOW QUALITY PROTEIN: dynein heavy chain domain-containing protein 1 [Orcinus orca] | 13% | 8.4 | 32% | XP_004279330.1 |
| SuperContig_1025868 | 1238 | DNA primase [Pseudaminobacter manganicus] | 23% | 9.2 | 32% | WP_080921700.1 |
| SuperContig_1035909 | 1185 | hypothetical protein T12_433 [Trichinella patagoniensis] | 16% | 5.5 | 31% | KRY11477.1 |
| SuperContig_1196200 | 734 | DUF948 domain-containing protein [Lactobacillus buchneri] | 41% | 4.7 | 27% | WP_014939867.1 |

The identification of a sex-linked *ATRX* homolog is notable as *ATRX* is known to play major roles in sex determination in mammals and other vertebrates (45–48). Alignments between scaffold SuperContig_990642 and the autosomal *ATRX* homolog revealed that two distinct *ATRX* homologs exist in axolotl (Figure 5). Alignments between *ATRX* and its sex-specific duplicate show polymorphisms in the *ATRX* gene that are not present in sex-linked *ATRX*, characteristic of a hemizygously-inherited duplication (Supplementary Figure 1). Henceforth, we refer to the conserved syntenic homolog on LG2 as *ATRX* and the W-specific homolog as *ATRW*. A nucleotide alignment between the axolotl *ATRX* and *ATRW* genes shows that the genes share 90% identity across 1089 aligned nucleotides, and as such it appears that the two genes diverged relatively recently by transposition of a duplicate gene copy to the W chromosome. To further test this idea and better define the timing of this duplication, several trees were generated using *ATRX* homologs from multiple vertebrate taxa (Figure 6, Supplementary Figure 2). Based on these trees, we infer that a duplication event gave rise to *ATRW* within *Ambystoma*, after divergence from its common ancestor with newt (the two lineages shared a common ancestor ~151 MYA) (49). Considering the degree of sequence divergence and the relative length of shared vs. independent branches we estimate that the *ATRW* homolog may have arisen sometime in the last 20 MY (Figure 6B), a timing that roughly coincides with a major adaptive radiation in the tiger salamander lineage (50, 51).

**Figure 5.**
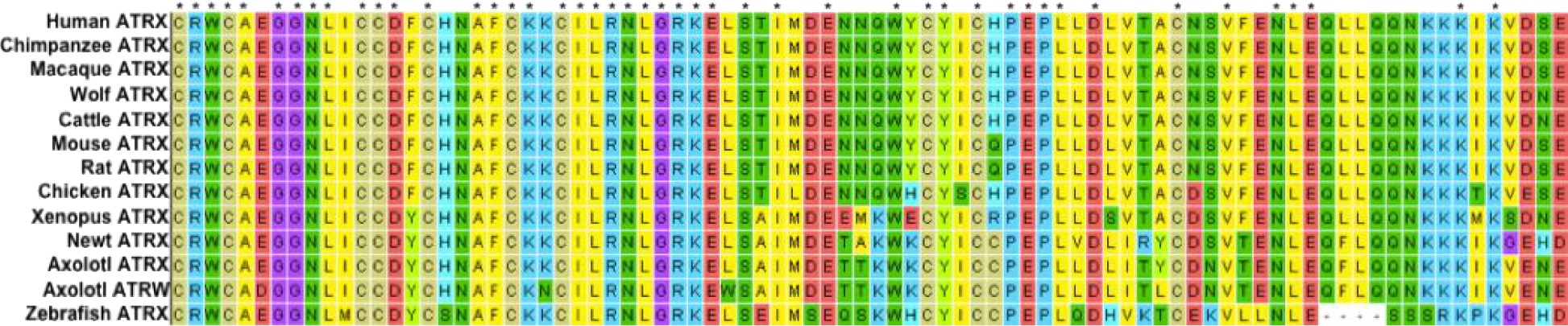
Alignment of translated nucleotides from *ATRX* in multiple vertebrate taxa. The alignment from MEGA7 (84) of 84 amino acids of *ATRX* with conservation in 12 vertebrates, including *ATRX* and *ATRW*from axolotl show the relative number of changes in codons specific to all amphibians, salamanders, axolotl and axolotl *ATRW*. A total of two out of nine nucleotide substitution events specific to the *ATRW* have altered the predicted codon.

To shed further light on the evolution of *ATRX* and *ATRW* within the *Ambystoma* lineage, we examined patterns of derived substitutions in *ATRX* and *ATRW*. Across the 251 bp alignment, 9 nucleotide substitutions can be attributed to *ATRW* since the divergence of axolotl, and these are associated with changes in 2 amino acids. By comparison, *ATRX* on LG2 shows only 1 nucleotide substitution since the duplication event (Figure 6). This suggests that *ATRW* may be evolving at a faster rate than *ATRX*, in which case 20 MY may represent a substantial overestimate for the origin of the duplication that gave rise to *ATRW*.

**Figure 6.**
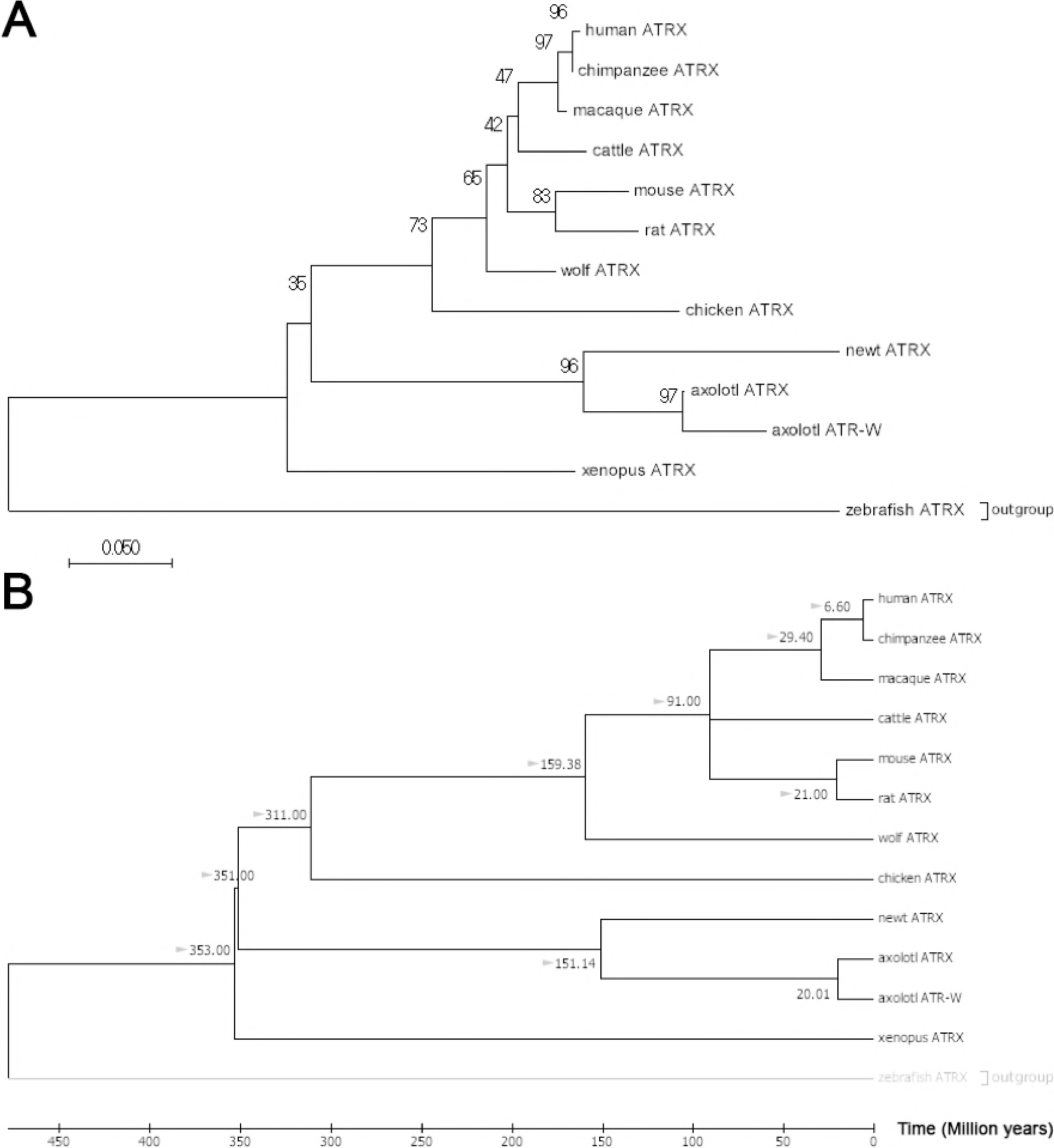
Neighbor-Joining trees for vertebrate *ATRX* with bootstrap and divergence time estimations. **A)** Evolutionary relationships among *ATRX* homologs were inferred using the Neighbor-Joining method (87). The bootstrap consensus tree inferred from 10000 replicates is taken to represent the evolutionary history of the taxa analyzed (88). The percentage of replicate trees in which the associated taxa clustered together in the bootstrap test (10000 replicates) are shown next to the branches (88). Evolutionary distances were computed using the Maximum Composite Likelihood method (89) and are in the units of the number of base substitutions per site. The analysis involved 13 nucleotide sequences. Codon positions included were 1st+2nd+3rd+Noncoding. All positions containing gaps and missing data were eliminated resulting in the inclusion of 251 positions in the final dataset. Evolutionary analyses were conducted in MEGA7 (84). **B)** A time-scaled phylogenetic tree inferred using the Reltime method (90) and estimates of branch lengths inferred using the Neighbor-Joining method as in A (87). The tree was computed using 10 calibration constraints. Divergence times estimated by Timetree were added manually and are marked with gray arrows (49). This tree indicates that the duplication event giving rise to *ATRW* in axolotl may have occurred ~20MYA.

## DISCUSSION

### Sex chromosome evolution in the axolotl

The results from this study show that the homomorphic sex chromosomes of the axolotl contain a small non-recombining region that is specific to the female W chromosome. The female-specific sequence is estimated to be approximately 300Kb, or roughly 1/100,000^th^ of the enormous axolotl genome. It is not surprising that the differences in recombination were not initially evident due to the physical size of the genome and marker density in the *Ambystoma* meiotic map (29). With respect to the current fragmented female genome assembly, it is still not possible to predict gene orders within this region or locate possible inversions; however, the data are sufficient to identify robust markers for sex and genes that exist in the non-recombining region. Of the few protein-coding genes found within the validated sex-specific scaffolds, two appear to represent non-repetitive coding sequences, including one that represents a relatively recent duplication of the transcriptional regulator *ATRX*.

The *ATRX* gene is located in the non-recombining region of the X chromosome in mammals. The gene encodes a chromatin remodeling protein that belongs to the SWI/SNF family. It is linked to the rare recessive disorder, alpha-thalassemia X-linked intellectual disability, which is characterized by severe intellectual disability, developmental delays, craniofacial abnormalities, and genital anomalies in humans. In some cases, a mutation in the *ATRX* gene can lead to female sex reversal due to early testicular failure (52, 53). Gene expression studies performed in a marsupial and eutherian showed that *ATRX* expression was highly conserved between the two mammals and was necessary for the development of both male and female gonads (48). Because *ATRX* is one of the few protein-coding genes present in the region of W-specific sequence and has been characterized in the sex differentiation of mammals, we propose *ATRW* as a candidate sex gene for axolotl, or alternately a strong candidate for an acquired, sexually antagonistic gene.

Reanalysis of expression data from recent published tissue-specific transcriptomes showed expression of the *ATRX* gene (from LG2) in all major tissues and developing embryos, however, they showed no evidence of expression of the *ATRW* gene (54). The tissues represented in the study included whole limb segments, blastemas from regenerating limbs, bone and cartilage, muscle, heart, blood vessel, gill, embryos, testis, and notably, ovaries. It is not clear at what stage the ovarian tissue was taken; however, the author suggests multiple ovaries were sequenced from an adult, and multiple libraries exist for the tissue. It is possible that this sex-specific gene is simply not highly expressed at this specific stage (or in the adult stage, in general) and may only be expressed during early gonadogenesis. Similarly, W-linked genes in chicken were unknown until RNAseq studies were performed prior to and during gonadogenesis (55).

If *ATRW* is the primary sex-determining gene in axolotl, then the origin of this gene marks the origin of sex chromosomes in the tiger salamander lineage. A time-scaled gene tree based on sequence substitution rates of *ATRX* genes in multiple vertebrate placed the *ATRX* duplication event at ~20 MYA (Supplementary Figure 1). This estimate places the *ATRX* duplication event within the *Ambystoma* clade but suggests that not all ambystomatids necessarily share the sex chromosome. Based on the *Ambystoma* species tree (49), we expect the same sex chromosomes and sex locus to be present in the tiger salamander species complex but not necessarily in the more distantly related *A. jeffersonianum* complex or deeper ambystomatid lineages (Figure 7).

**Figure 7.**
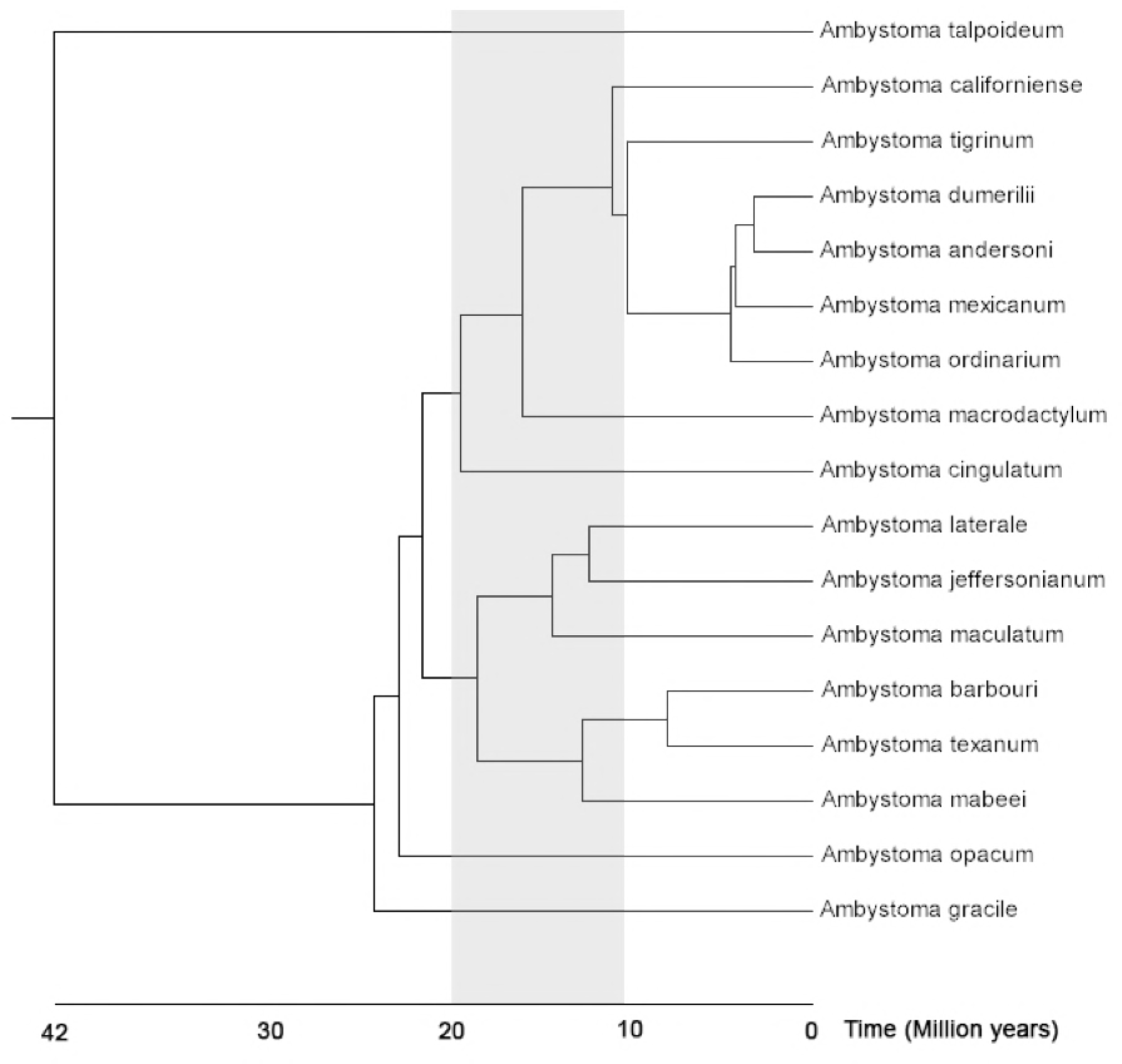
A species tree for the genus *Ambystoma*. The gray shaded region shows the approximate timing of the *ATRW* duplication event. The tiger salamander complex consists of 7 named species that occur in the same monophyletic clade as *A. californiense, A. mexicanum*, and *A. tigrinum* (56, 91). This tree was generated using Timetree (49) with modification to the position of *A. californiense* based on Shaffer and McKnight (1996) and Shaffer et al. (2004).

Given the relatively recent origin of *ATRW*, species within the tiger salamander complex are predicted to contain the same sex chromosomes. The tiger salamander species complex consists of more than 30 named species that encompass a range of diversification dates (50, 56). Further analyses of sex determination within this complex should therefore facilitate future studies aimed at more precisely characterizing the timing of the *ATRX/W* duplication and the evolution of other W-specific sequences. Ongoing improvements to the *Ambystoma* genome assembly and development of genome assemblies for other salamander taxa should improve our ability to assess hypotheses related to the presence of homomorphic sex chromosomes (*e.g*. recent evolution, high-turnover, and fountain of youth) (1, 17, 57–62). Additionally, recent efforts to develop genetic tools for the axolotl model should facilitate functional analyses that will be necessary to test whether *ATRW* is the primary sex-determining gene in axolotl or elucidate its role as a sexually antagonistic factor (63, 64). Methods for achieving targeted gene knockout and knock-ins have been developed in axolotl and could be adapted to better assess the functionality of *ATRW* in axolotls (40, 65, 66).

### Utility of sex-linked markers in axolotl research

Sex is an important biological variable in research, as it may contribute to variation in experimental studies. Because axolotl is an important model for many areas of research and has shown sex-specific effects, such as tail regeneration, it is important for investigators to differentiate sex effects from other experimental variables (28). Until now it was necessary to visualize the sex organs, utilize axolotls that had produced gametes, or perform experiments in hybrid crosses that segregate markers at the linked locus E24C3 in order to accurately determine sex in axolotls (29). However, many experiments utilize juvenile animals that may not have completed gonadal differentiation or maturation. With several robust markers for W-specific sequences in hand, it is now possible to precisely differentiate sex of an axolotl with a simple PCR (67). These markers will also positively impact axolotl husbandry, as individuals may be housed and utilized in experiments accordingly.

## MATERIALS AND METHODS

### Laser capture microdissection and amplification

Preparation of cells for metaphase spreads and laser capture were performed as previously described (30). Briefly, fixed cells were spread on UV-treated 1.0mm polyethylene naphthalate (PEN) membrane slides. Slides were inverted (membrane side down) over a steam bath of distilled water for 7 seconds. Immediately after steaming, 100 μl of the fixed cells were dropped across the middle of the slide lengthwise. Each slide was subsequently placed in a steam chamber at ~35°C for 1 minute, then set on the hot plate for 5 minutes. After slides dried, chromosomes were stained via immersion in freshly made Giemsa stain (Sigma-Aldrich GS500-500 ML: 0.4% Giemsa, 0.7 g/L KH2PO4, 1.0 g/L Na2HPO4) for 2 minutes, rinsed in 95% ethanol, rinsed in distilled water, then allowed to dry in a desiccator until used.

The sex chromosome was captured using a Zeiss PALM Laser Microbeam Microscope at 40X magnification as previously described (30). The sex chromosome was dissected individually using a Zeiss PALM Laser Microbeam Microscope at 40X magnification and catapulted into a Zeiss adhesive cap tube (Zeiss 415190-9191-000). 10 μl of a chromatin digestion buffer was pipetted into the cap (30) and the tube was kept inverted overnight at 55°C. After incubation, the sample was centrifuged briefly and incubated at 75°C for 10 minutes and 95°C for 4 minutes to inactivate the Proteinase K. Along with 23 other samples, the sex chromosome sample was immediately carried through full amplification using the Rubicon PicoPlex DNAseq Whole Genome Amplification (WGA) kit (R30050). Amplification followed the standard manufacturer protocol, with one exception: a chromatin digestion step replaced the cell extraction step. After amplification, an Agilent 2100 Bioanalyzer and accompanying DNA 12000 kit (Agilent DNA 12000 Kit 5067-1508) was used to estimate concentration and size distribution. The sex chromosome sample had a concentration >9ng/μl and was sequenced on an Illumina HiSeq 2500 (Hudson Alpha Institute for Biotechnology, Huntsville, Al). After initial sequencing, the same sample was further sequenced to generate paired-end 150bp reads on a full lane of HiSeq 2500.

### Sequence analyses and assembly

Because amplified sequences contain a non-complex leader sequence corresponding to the pseudorandom primers that are used for whole chromosome amplification, reads were trimmed prior to further processing. Trimmomatic was used to remove leader sequences derived from phiX and to trim any window of 40 nucleotides with quality score lower than Q30 (68). Reads were then aligned to 945 model transcripts from the *Ambystoma* linkage map (35) using the Burrows Wheeler Aligner with the single-end mapping option and BWA-MEM algorithm (69). They were also aligned to several bacterial genomes as well as the human reference genome using the paired-end mapping option to identify exact matches for Bowtie 2 (70). Paired reads that mapped concordantly to the human and bacterial genomes were considered potential contaminants and removed. After trimming and removal of potential contaminants, the reads were corrected with Blue (71) using female *A. mexicanum* whole genome shotgun data (30) and assembled with SOAPdenovo2 (72).

To assign scaffolds from the whole genome assembly of a male axolotl genome to the Z chromosome, error-corrected laser capture reads were aligned as paired-end reads to the assembly with BWA-MEM and filtered to preserve only pairs with concordant reads that map to the reference with no mismatches (69). For each scaffold we calculated physical coverage (i.e. coverage by paired-end fragments: bedtools v. 2.27, genomeGoverageBed, option pc, (73)) and assigned scaffolds to the Z chromosome if at least 5% of their bases were covered by reads from laser capture sequencing.

### FISH of sex-associated BAC *E24C3*

Fluorescent *in situ* hybridization of BACs to metaphase chromosome spreads were performed as previously described (74, 75). A Qiagen Large Construct kit (Qiagen Science, 12462) was used to extract bacterial artificial chromosome (BAC) DNA for *E24C3* and *E12A6*, previously associated with sex (29). Probes for *in situ* hybridization were labeled by nick-translation using direct fluorophores Cyanine 3-dUTP (Enzo Life Sciences, ENZ-42501) or Fluorescein-12-dUTP (Thermo Scientific, R0101) as described previously (74) and hybridization of BAC probes was performed as previously described for axolotl chromosomes (40).

Phenol-chloroform extraction in 1.2X SSC was used to isolate repetitive DNA fractions from female salamander tissue (76). DNA was denatured for 5 minutes at 120°C, re-associated at 60°C for 1 hour to obtain C_o_t DNA. Microtubes containing the DNA were placed on ice for 2 minutes, then transferred to a bead bath at 42°C for 1 hour with 5X S1 nuclease buffer and S1 nuclease for a concentration of 100 units per 1 mg DNA. DNA was precipitated with 0.1 volume of 3M sodium acetate and 1 volume isopropanol at room temperature, tubes were inverted several times and centrifuged at 14,000 rpm for 20 minutes at 4°C. DNA was washed with 70% ethanol, centrifuged at 14,000 rpm for 10 minutes at 4°C, air dried and solubilized in TE buffer.

### Conservation and evolution of salamander chromosomes

To evaluate the sex chromosome assembly, we performed alignments between the sex chromosome assembly and reference transcripts (V4: Sal-Site)(32) using megablast (77) to identify genes that occur on the sex chromosome. These genes were then aligned (tblastx) (78) to annotated protein coding genes from the chicken genome assembly (Gallus_gallus-4.0). Annotated genes from scaffolds assigned on the basis of read mapping were aligned (blastp) (78) to this set of annotated chicken genes. Those with an alignment length of at least 50 amino acids and at least 60% identity were considered potential homologs.

A similar approach was taken to identify the homologous newt linkage group to assess potential sex candidate genes. *Ambystoma* reference transcripts from LG9 (V4) were aligned (tblastx) (78) to the chicken genome assembly (41). Using the same minimum thresholds as above, the potential homologs were then used to blast (tblastx) (78) to the newt, *Notophthalmus viridescens*, reference transcripts (36).

### Identification of female-specific regions

We applied depth of coverage analysis to identify single-copy regions in the assembly that have approximately half of the modal coverage in females and underrepresented/absent coverage in males. Reads were generated on an Illumina HiSeq2000 (Hudson Alpha Institute for Biotechnology, Huntsville, Al.) from DNA that was isolated via phenol-chloroform extraction (76) from 48 individuals that were drawn from a previously described backcross mapping panel (42). The resulting reads were aligned to the axolotl draft genome assembly using BWA-MEM (using default parameters) followed by filtering of secondary alignments (samtools view – F2308) and alignments clipped on both sides of the read. Merging of female and male bam files was performed using *Samtools merge* (69, 79).

We used DifCover (https://github.com/timnat/DifCover) (80) to identify candidate female-specific regions. The method works by computing the ratio of female:male average depth of coverage across continuous intervals containing approximately *V*, valid bases. Valid bases are defined by lower and upper limits on depth of coverage for females (f) and males (m), respectively designated as min*f*, min*m*, max*f* and max*m*. If *Cf* and *Cm* define female and male coverage for a given valid base, then 1) *Cf* < max*f and Cm* < max*m*; and 2) *Cf*>min*f or Cm*>min*m*. The upper limits identify and allow skipping of fragments that contain repeats, while the lower limits serve to exclude underrepresented fragments with small numbers of reads in both males and females.

After testing, we chose V=1000 and assigned lower limits equal to one third of modal coverage, (8 for females and 9 for males) and upper limits 3X of modal coverage, (75 for females and 87 for males). The enrichment scores [log2(standardized sperm coverage/blood coverage)] were computed for each interval. If the average coverage in males for an interval was zero, we replaced the coverage estimate with a non-zero positive value corresponding to alignment of half of one read. Some intervals were shorter than 1Kb and contained fewer than 1000 valid bases (short scaffolds or intervals that fall on the scaffold ends). These shorter intervals were filtered to exclude intervals with fewer than 500 bases or fewer than 200 valid bases.

Scaffolds that were validated through PCR in a panel of 6 females and 6 males were aligned to the V4 and V5 *Ambystoma* transcriptome assemblies in order to identify the genes present on the W-specific portion of the sex chromosome. If a transcript aligned to the scaffold with a percent identity higher than 95%, that transcript was blasted (blastx) (78) to the NCBI nonredundant protein database to search for homologous genes.

### Primer design and PCR

Primers were designed within the sex candidate regions identified using Primer3 (81). Each primer was 25-28 bp in length, with a target melting temperature of 60°C, 20-80% GC content and 150-400 bp product sizes depending on the size of the region and location of repeats (avoiding inclusion of repetitive sequence in primer and product). Fragments were amplified using standard PCR conditions (150ng DNA, 50 ng of each primer, 200 mM each dATP, dCTP, dGTP, dTTP; thermal cycling at 94°C for 4 minutes; 34 cycles of 94°C for 45 seconds, 55°C for 45 seconds, 72°C for 30 seconds; and 72°C for 7 minutes). Reactions were tested on a panel of six males and six females to validate sex specificity. Gel electrophoresis was performed and presence/absence was recorded for each set of primers (Supplementary Figure 3). The scaffolds from which primers were designed were considered female-specific if the primers yielded specific amplicons in all six females and in no males.

Results from these data were used to develop a PCR based assay for determining sex in axolotls at any stage of development. This method uses a primer pair that amplifies a 219 bp DNA fragment in females and an internal control that yields a 486 bp DNA fragment in both sexes. This biplex PCR results in two bands (219 bp and 486 bp) for females and only the control band (486 bp) in males (67).

### Phylogenetic Reconstruction

Homologene was used to collect putative homology groups from the *ATRX* genes in a variety of eukaryotes (82). Sequence for axolotl *ATRX* was obtained from *Ambystoma* reference transcripts, and the newt *ATRX* gene was obtained by aligning human *ATRX* to the newt reference transcriptome (83). All sequences were aligned using MEGA7 (84) via MUSCLE (85). Sequences were trimmed to compare a conserved subregion of the sequence that was present in all species, a string of 251 codons (Figure 5). Divergence time estimates were drawn from the TimeTree webserver (49).

### Ethics Statement

All methods related to animal use were performed in accordance with AAALAC guidelines and regulations, under supervision of Division of Laboratory Animal Resources. Tissue collection was performed in accordance with protocol number 01087L2006, which was approved by the University of Kentucky Office of Research Integrity and Institutional Animal Care and Use Committee.

**Supplementary Figure 1.**
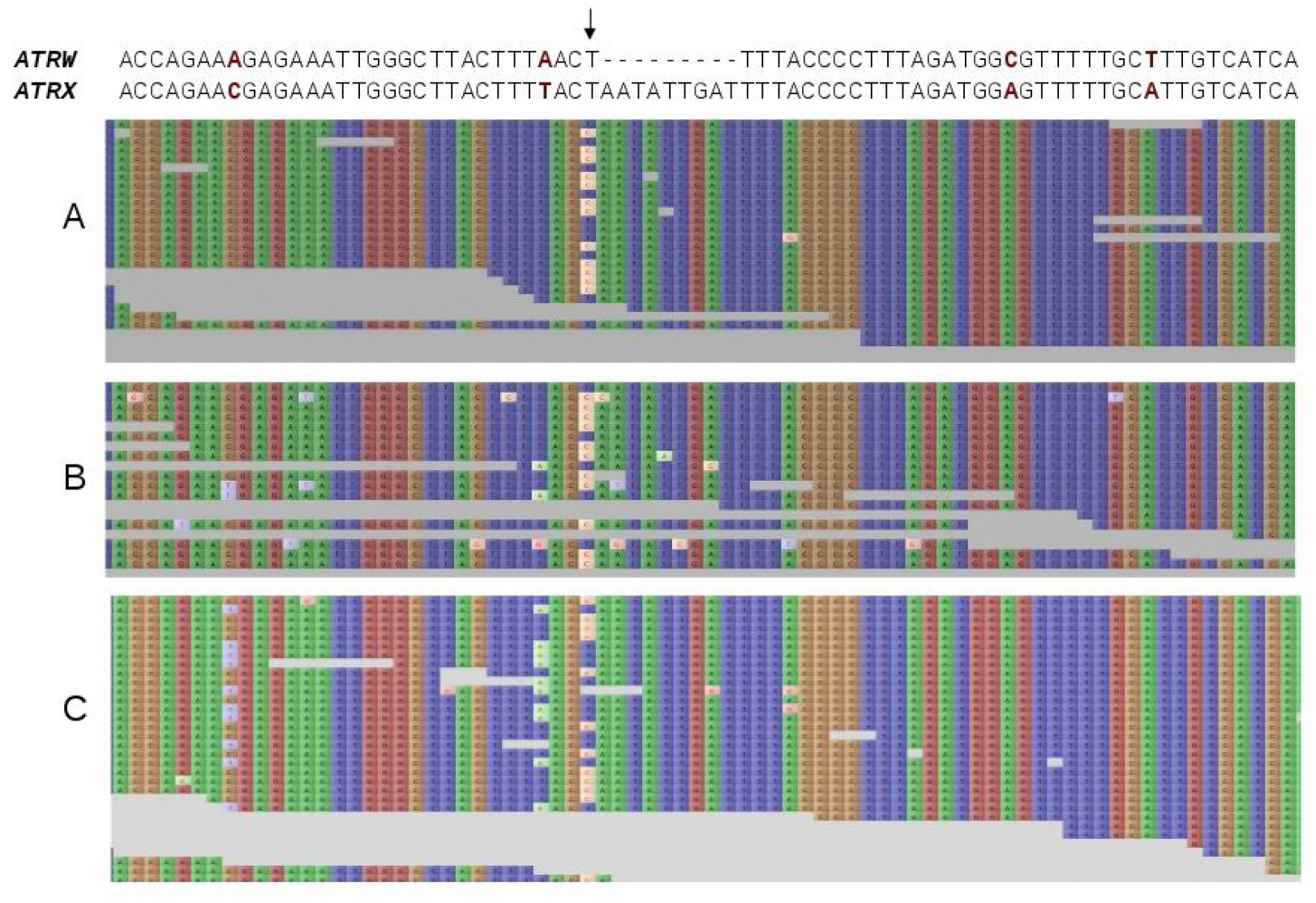
Two alleles detected for *ATRX* in axolotl. Two alleles can be observed in exon 9 of ATRX within the region homologous to the validated sex-specific DNA fragment of *ATRW* (SuperContig_990642). Alignment of short reads from genome sequencing of **A)** an *A. mexicanum* female, **B)** 22 female individuals, **C)** and 26 males to positions 6590-6665 of SuperContig_546209 (the contig containing exon 9 of the ATRX gene in the female genome assembly). The position marked with an arrow corresponds to position 1018 of SuperContig_990642 and position 6620 of SuperContig_546209. Visualization of the alignment produced using Tablet 1.14 (92).

**Supplementary Figure 2.**
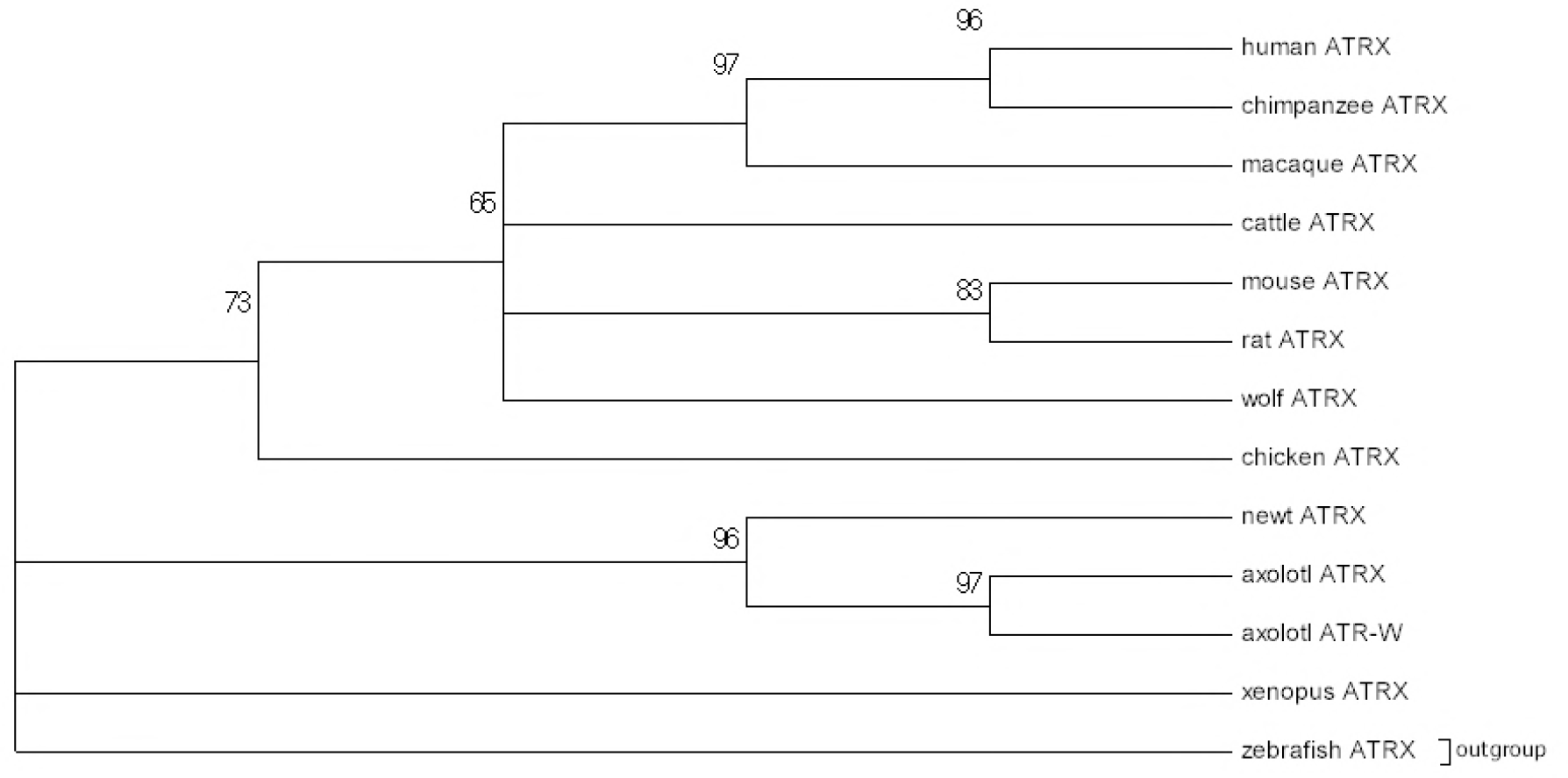
Neighbor-Joining tree with bootstraps for vertebrate *ATRX*. The evolutionary history was inferred using the Neighbor-Joining method (87). The bootstrap consensus tree inferred from 10000 replicates is taken to represent the evolutionary history of the taxa analyzed (88). Branches corresponding to partitions reproduced in less than 50% bootstrap replicates are collapsed. The percentage of replicate trees in which the associated taxa clustered together in the bootstrap test (10000 replicates) are shown next to the branches (88). The evolutionary distances were computed using the Maximum Composite Likelihood method (89) and are in the units of the number of base substitutions per site. The analysis involved 13 nucleotide sequences. Codon positions included were 1st+2nd+3rd+Noncoding. All positions containing gaps and missing data were eliminated. There was a total of 251 positions in the final dataset. Evolutionary analyses were conducted in MEGA7 (84).

**Supplementary Figure 3.**
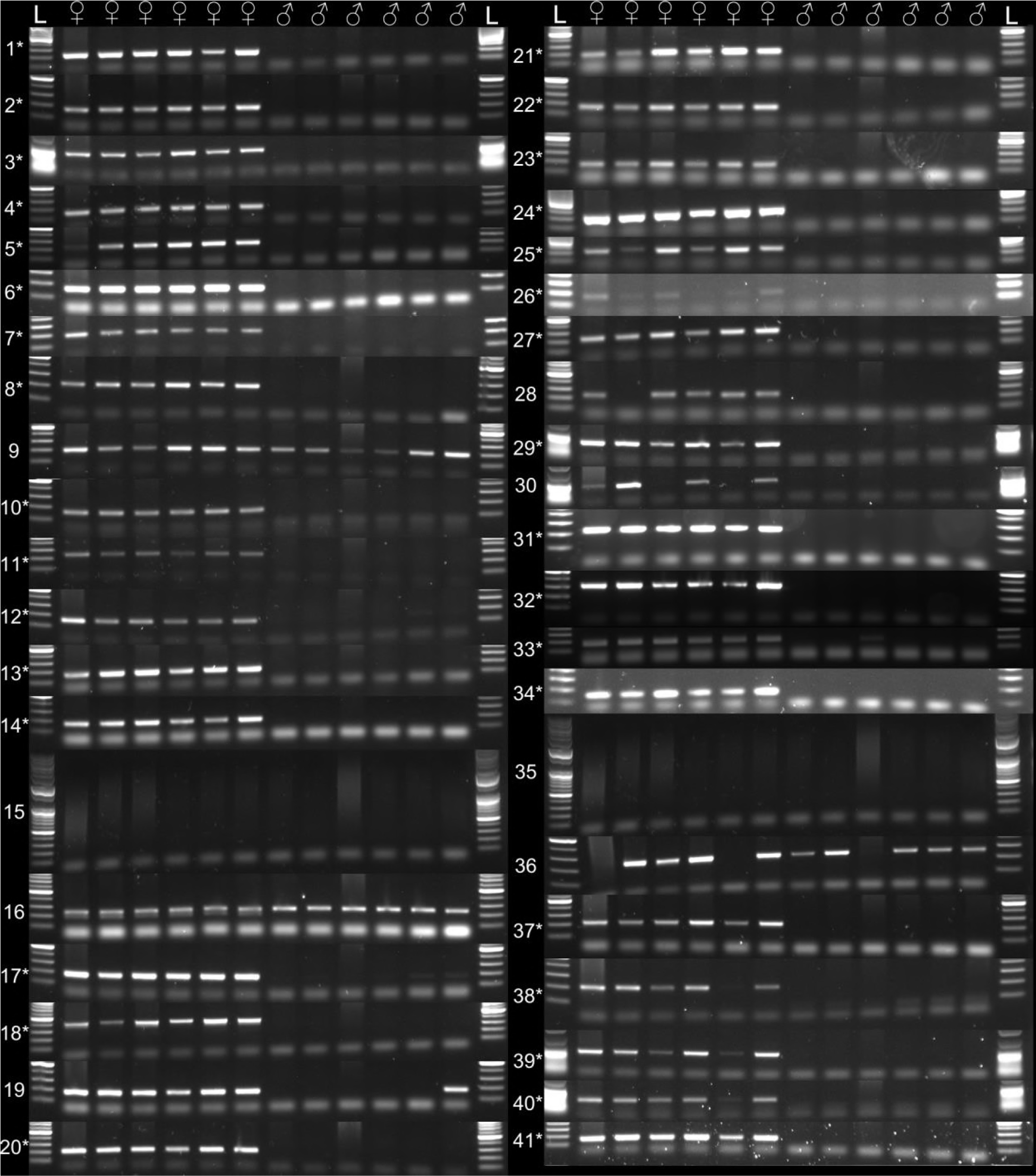
PCR validation of sex-specific fragments. Gel electrophoresis of PCR for scaffolds determined to be sex-specific based on computational analyses. Those that show presence in females only are denoted with an asterisk and considered sex-specific. PCRs were tested on six females and six males, and the associated lanes are denoted with ♀ and ♂, respectively. The first and last lanes are labeled with “L” to denote 100 bp ladder. Numerical labels correspond to primer information provided in Supplementary Table 3.

**Supplementary Table 1**- Coverage statistics for scaffolds that were assigned to the *Ambystoma* Z chromosome. Scaffolds identifiers correspond to the published assembly

**Supplementary Table 2**- Alignment statistics and estimated coverage ratios for regions that show enrichment in females.

**Supplementary Table 3**- Summary of PCR assays for predicted female-specific regions.

